# AbFold -- an AlphaFold Based Transfer Learning Model for Accurate Antibody Structure Prediction

**DOI:** 10.1101/2023.04.20.537598

**Authors:** Chao Peng, Zelong Wang, Peize Zhao, Weifeng Ge, Charles Huang

## Abstract

**Motivation:** Antibodies are a group of proteins generated by B cells, which are crucial for the immune system. The importance of antibodies is ever-growing in pharmaceutics and biotherapeutics. Despite recent advancements pioneered by AlphaFold in general protein 3D structure prediction, accurate structure prediction of antibodies still lags behind, primarily due to the difficulty in modeling the Complementarity-determining regions (CDRs), especially the most variable CDR-H3 loop.

**Results:** This paper presents AbFold, a transfer learning antibody structure prediction model with 3D point cloud refinement and unsupervised learning techniques. AbFold consistently produces state-of-the-art results on the prediction accuracy of the six CDR loops. The predictions of AbFold achieve an average RMSD of 1.51 Å for both heavy and light chains and an average RMSD of 3.04 Å for CDR-H3, bettering current models AlphaFold and IgFold. AbFold will contribute to antibody structure prediction and design processes.

## Introduction

Antibody, also known as immunoglobulin, is a special class of protein. Antibodies are produced by B cells when the adaptive immune system responds to invading antigens. Antibodies can recognize and bind to antigens with high specificity and affinity. Moreover, antibody-based drugs have recently become the fastest-growing biological pharmaceutics, especially in the field of anticancer, autoimmune, and vaccine drugs[1, 2]. However, the study and design of antibodies still face significant challenges. The most challenging part is to accurately predict antibody 3D structure, which determines antibody-antigen binding interactions such as affinity and specificity. Thus, understanding antibody function based on its 3D structure has become ever more important from both industry and academic perspectives.

A Y-shaped antibody has two symmetric parts, each composed of a heavy chain and a light chain. Both heavy and light chains consist of one variable domain and a few constant domains. While the structures of the constant domains are highly conserved within a species, the structures of the variable domains are diverse. The variable domain (F_v_) regions are situated at the tip of Y-shaped antibodies and determine the functionality of antibodies. Inside the F_v_, there exit six hypervariable regions, known as complementarity-determining regions (CDRs). They are three loops for the heavy chain, CDR-H1, H2, and H3, and three loops for the light chain, CDR-L1, L2, and L3. The six CDR regions are the determining factor for the antibody-antigen interactions, thus, are the primary targets for structural modeling. CDR-H3 situates at a close region between the heavy chain and light chain, and its conformation is affected by the inter-chain orientation, adding complexity to the structure prediction [3, 4]. Hence, CDR-H3 structure prediction has been the focal point of recent antibody structure research [4].

Since most components of the F_v_ tend to adopt canonical folding structures, grafting prediction methods are able to reach an average root-mean-square deviation (RMSD) of less than 1 Å, based on the previously determined structure with similar sequences [5-8]. However, these grafting methods do not predict the CDR-H3 loop structure well due to the high variability in sequence[9]. Recently, deep learning methods have dramatically improved the accuracy of general protein structure prediction. Moreover, deep learning methods have been demonstrated to outperform traditional experimental methods at a fraction of the costs and time [10, 11]. AlphaFold [12] can produce admirable prediction results on par or better out of the box without any optimization for the antibody sub-domain. Furthermore, AlphaFold-Multimer [13] continues pushing the prediction accuracy of protein complexes. In the area of antibody protein structure prediction, a few deep-learning models have been proposed, and the state-of-the-art method IgFold has achieved an RMSD of about 3 Å for the H3 loop [14]. The neural network architecture of IgFold is a fast end-to-end deep learning model utilizing a pre-trained BERT language model [15], a simplified AlphaFold-like transformer, and an invariant point attention structure. DeepAb [16] consists of a deep residual convolutional network for predicting inter-residue distances and orientations and a Rosetta-based protocol for generating structures from network predictions. ABlooper [17] predicts CDR loops with an end-to-end adapted graph neural network (GNN) architecture.

Although state-of-the-art deep learning computational methods have been shown to surpass the experimental ones on general protein structure prediction and have also produced promising results on antibody structure prediction [12, 16], they still face significant challenges in predicting the structure of CDR loops accurately. First, the experimental ground truth structure data of about 1,700[16] is quite limited to building a machine learning model. We try to overcome this data limitation with dataset augmentation methods. Second, CDR loops and the rest of the Fv parts adopt quite different conformations, and present a challenge for any single model to predict both well simultaneously. Therefore, we propose developing a new model specifically for CDR loop structure prediction.

In this paper, we present a new state-of-the-art model AbFold: a transfer learning model for the antibody structure prediction, by harnessing the prowess of generic protein structure model AlphaFold, and optimizing further with the point cloud model, producing more accurate predictions of an overall RMSD 1.51 Å, especially an improvement of 7% for the H3 loop. AbFold also exploits the self-supervising technique to obtain a much larger high-confidence training dataset, enabling the model to learn a more comprehensive knowledge of CDR loop conformation. With innovative neural network architecture and better CDR-H3 prediction capability, AbFold contributes to the state-of-the-art design process of novel therapeutic antibodies. All the code and dataset of this paper and pre-trained model will be released.

## Results

### Methodology overview

AbFold is a transfer learning model for accurate antibody structure prediction (Figure 1). The model has two inputs, the first is the heavy and light chain sequences of the antibody, and the second is the preprocessed backbone feature of the same heavy and light chain sequences. Single and pair representations provide insights into the input sequences and are then combined with the 3D point cloud-optimized backbone feature. And finally, the designated variational autoencoder (VAE) predicts the refined prediction structure.

**Figure 1.**
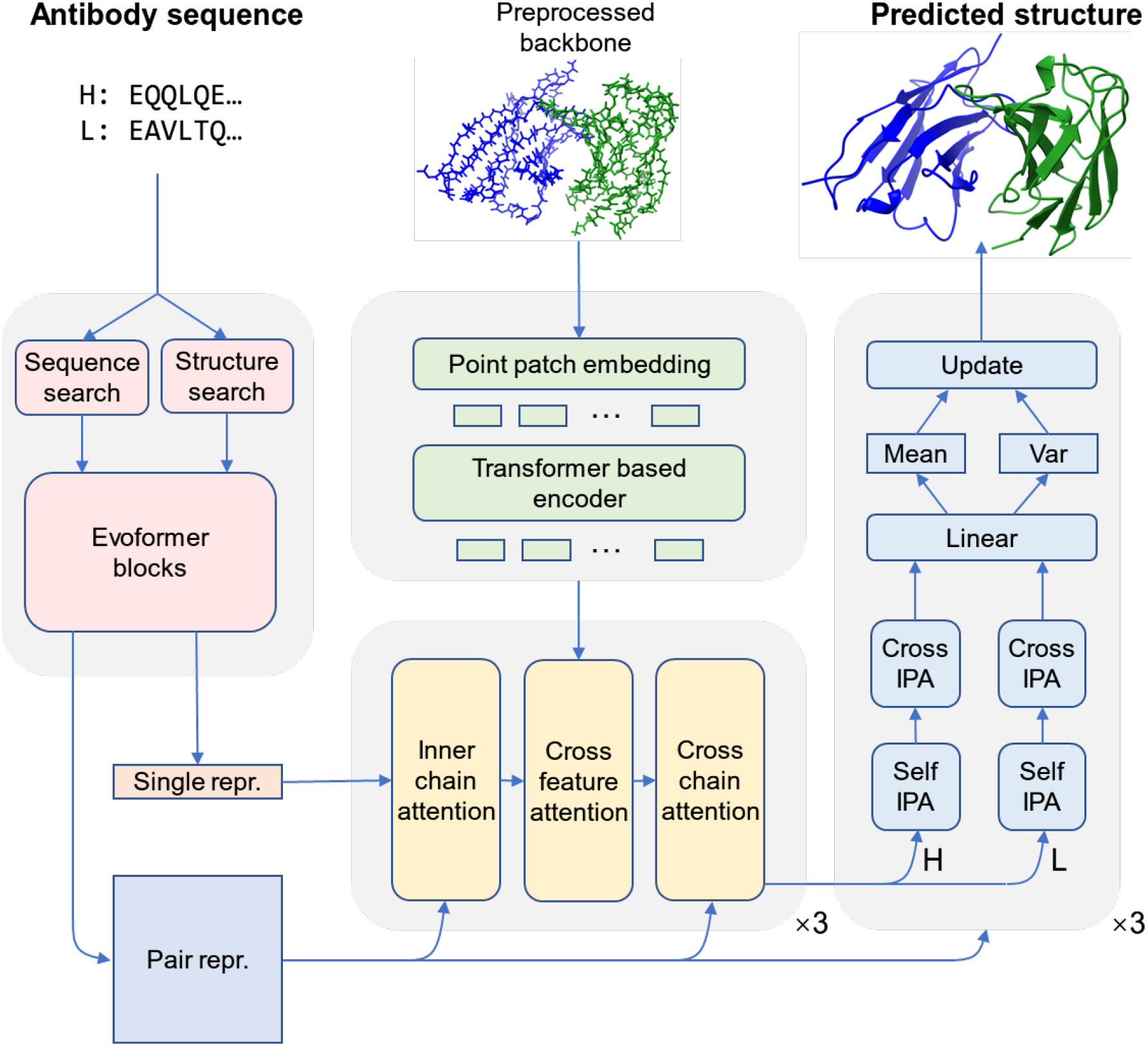
Model Structure of AbFold. Arrows show the information flow among the components described in this paper. Antibody sequences are converted to single and pair representations by Evoformer of AlphaFold2. Single representations are then updated by point cloud information using a series of attention blocks. Single and pair representations are used to directly predict atomic coordinates in later structures.

### Self-supervised learning for dataset augmentation

Training an accurate model for antibody structure prediction is challenging due to the limited number of experimentally tested antibodies, which stands at only a few thousand according to SabDab [18]. Methods have been proposed in prior works to deal with this data limitation problem, for example, in IgFold [14], a pre-trained language model extracts patterns from millions of natural antibody sequences to get the evolutionary embeddings; and in DeepAb [16], a recurrent neural network (RNN) encoder-decoder model obtains evolutionary and structural representations of antibody sequences. We diverge from existing methods since the representations they provided do not have enough information for determined CDR H3 prediction. AbFold utilizes the latest student-teacher self-supervised learning techniques[27], designed to capture general patterns from the sequences that may not be readily evident in the smaller subset of sequences with known structures and to embed them into a latent representation. Both teacher and student networks have the same architecture, but different parameter updating methods. The student network updates parameters through backpropagating gradients while the parameters of the teacher network are updated by the exponential moving average of the student network parameters. The student-teacher model is trained using a combination of sequences with and without structure. These two kinds of data simultaneously exist in a training batch and are put into both teacher and student networks in parallel. These sequences with experimental structures are used as supervised data for the student model directly. While those sequences without structures are first fed to the teacher model to get structure, predicted structures are used as supervised data for the student model.

### Transfer learning with antibody domain-focused architecture / Feature extraction

Our antibody structure prediction model, AbFold, transfers the general protein structure prediction model [12] to the antibody sub-domain by transferring learned knowledge of the Evofomer block.

AlphaFold, being a powerful generic deep learning model in the large field of protein structure prediction, is ideally suited to be fine-tuned to predict the antibody sub-domain structure with a limited amount of labeled structure data (only a few thousand) and more unlabeled data (hundreds of millions).

Transfer learning can transfer the knowledge learned from the source domain to the target domain and solve problems in the target domain despite a lack of labeled data. In the task of antibody structure prediction, there are few antibodies whose structure is known, so it is difficult for a neural network to learn the structural characteristics of antibodies from a small amount of labeled data. Therefore, this paper uses the AlphaFold2 model, trained on massive protein structures, to extract the basic features of antibody structures. The features extracted from AlphaFold2 are used for antibody structure prediction in subsequent processing.

Specifically, AlphaFold2-Multimer was selected to extract features. AlphaFold2-Multimer provides the top five groups of model parameters, from which we simply select the first model to extract features in AbFold. To overcome the long-time to search for homologous sequences in AlphaFold2 (around one hour per antibody), we used ColabFold[28] which takes a shorter time to complete the homologous sequence search. In the process of feature extraction, the input is the paired amino acid sequences of heavy and light chains, and the output is the Single representation and Pair representation output of the Evoformer module after the third Recycle of AlphaFold2-Multimer. The dimensions of Single representation and Pair representation are N × 384 and N × N × 128, respectively, where N is the sum of the lengths of the heavy and light chains.

### Point cloud for representation optimization

We employed a modified and pre-trained Point-MAE [19] to transform structures predicted by IgFold into features, which were then combined with features produced by the Evoformer module.

We modified the input part of the Point-MAE network to accommodate the data structure of antibodies. First, for atoms in proteins, we recognize that position information alone is insufficient, and the most critical property of an atom is its type, so we convert the structure of the antibody to 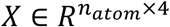, where *n*_*atom*_ denotes the number of atoms of the antibody and 4 denotes features of an atom (*x, y, z, automtybe*). Second, in our work, patches are divided based on *Cα* of amino acids. Every *Cα* is chosen as the center point of a patch representing the corresponding amino acid, then the K-nearest neighbor method is used to sample points around the *Cα* to form one patch. The patch size is fixed at 32 to ensure every patch can cover one complete amino acid. Furthermore, we crop antibody data of various lengths into sub-parts of fixed length for batch training. After sampling and cropping, a batch of antibody data with batch size *b*, patch number *n*, and patch size *k* is denoted as *P* ∈ *R*^*b*×*n*×*k*×4^.

Pre-training of the Point-MAE module is a self-learning process, which involves of masking out and predicting some patch positions with information from the left-out unmasked patches. A masked patch is replaced with a share-weighted learnable token, while an unmasked patch token is generated by a lightweight PointNet [20]. In addition, we utilize a multilayer perceptron to generate a position embedding for each patch. Both the encoder and decoder of this module consist of the transformer blocks. Moreover, every transformer block takes position embeddings as part of its input. The encoder exclusively handles unmasked tokens and produces embedded tokens as output, while the decoder takes both embedded tokens and masked tokens as input, and generates the decoded masked tokens as output. These tokens are further processed by the prediction head to get predicted atom positions. The Point-MAE model was pre-trained using a dataset of 92,300 unlabeled antibodies whose structures are computationally predicted with IgFold.

### VAE for variable conformations of CDR loops

Recent studies have demonstrated that the CDR loops of antibodies exhibit multiple conformations, indicating that their structures are dynamic and may change based on antigen and environmental factors [21]. Therefore, in this paper, we propose to use a variational autoencoder (VAE) to model the dynamic nature of antibody structures, over traditional fixed network models.

For a batch of sequence feature 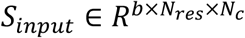, a linear layer is applied to transform it to 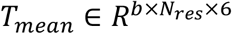 and 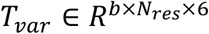 where *T*_*mean*_ and *T*_*var*_ separately stands for mean value and variance value of (*x, y, z*) in the translation matrix and (*b, c, d*) in the rotation matrix. In the update stage, we re-parameterize Euclidean transform[12] by *ΔT* = *mean* + *x ⋅ var*, where *x* is sampled from *N*(0, 1) and 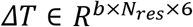. Lastly, we apply the Euclidean transform to earlier transform to get the updated structure: *T* = *T*∘ *ΔT*.

### Fusing information to improve CDR H3 accuracy

CDR H3, which refers to the third complementarity-determining region of the heavy chain, is known to be the most variable part of the antibody structure. It can vary in length from 5 to 26, and CDR H3 with a long loop often exhibits very low sequence similarity, making it difficult to predict using a limited set of conformations [25]. As a result, deep learning methods for predicting antibody structures struggle to accurately predict the atomic coordinates of CDR H3 compared to other CDR loops. Among existing methods, IgFold and AlphaFold-Multimer are the two best models for predicting antibody structures. However, they often predict distinct conformations of CDR H3 loops, with one model predicting a more accurate conformation than the other. To improve the accuracy of predicted antibody structures in CDR H3, we combine the strengths of both methods. Specifically, we use AlphaFold-Multimer to generate sequence embeddings and Point-MAE to extract point cloud features from the IgFold-predicted structure. We then merge the sequence embeddings and point cloud features using cross-attention.

### Datasets

We obtained paired antibody structures from The Structural Antibody Database (SAbDab) and applied strict filters to get structures with sequence similarity ≤99% and resolution ≤4.0 Å. In order to make a meaningful comparison with models which updates and utilizes the latest data, such as IgFold, we divide our dataset into the training set and the test set based on the date of structure deposition. Specifically, structures deposited before April 24, 2022 were used for training, while those deposited after that date were used for testing. Overall, our supervised training dataset comprises 1847 paired structures, while our test dataset includes 24 paired structures. In addition, we used 20,000 heavy-and-light-chain paired antibody sequences from the Observed Antibody Space (OAS) database [26] as unsupervised input. Within each mini-batch of the training dataset, supervised data and unsupervised data were divided equally.

### Training procedure

The model is trained with three types of loss functions, including Frame Aligned Point Error (FAPE) loss, structural violation loss, and bond length loss. The total loss function is defined as follows:

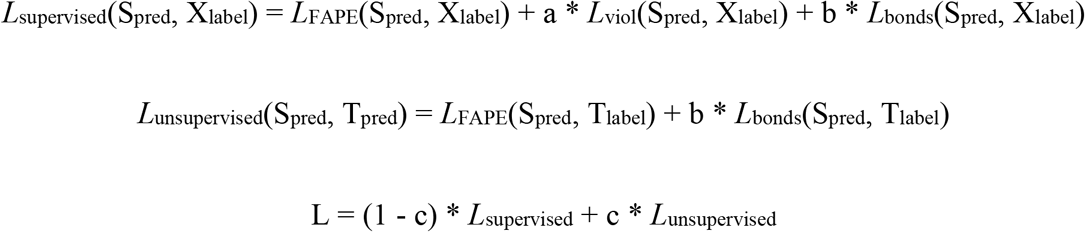

where S_pred_, X_label_, and T_pred_ represent the predicted structure of the student network, the experimental structure, and the predicted structure of the teacher network, respectively. The training process consists of two stages: the initial training stage and the fine-tuning stage. In the initial training stage, the model is trained for 150 epochs with a=0, b=0, and c=0.5. Subsequently, in the fine-tuning stage, the model is trained for 150 epochs with a=1, b=1, and c=0.1.

For optimization, we use an AdamW optimizer with a base learning rate of 10^−4^. The learning rate linearly increases from 0 to 10^−4^ in the first 1000 iterations and further decreases by a factor of 0.95 every 1000 iterations after 10000 iterations. The size of the mini-batch is set to 32. To avoid exploding gradient problem, we apply gradient clipping by 2-norm on all parameters, with a clipping value of 0.1.

### AbFold outperforms other methods in CDR H3

We compared the performance of AbFold with other deep-learning methods for F_V_ structure prediction. For every antibody in the benchmark, we predicted its structure with AlphaFold-Multimer, ImmuneBuilder, DeepAb, EquiFold, IgFold, and AbFold. The most challenging aspect of antibody structure prediction is modeling the CDR loops. So we measured the backbone heavy-atom RMSD between the predicted structure and the native structure of each region in Fv (including framework region and CDR loops) with the Chothia scheme-defined CDR loops. Table 2 provides a detailed breakdown of the results for each method.

**Table 2:**
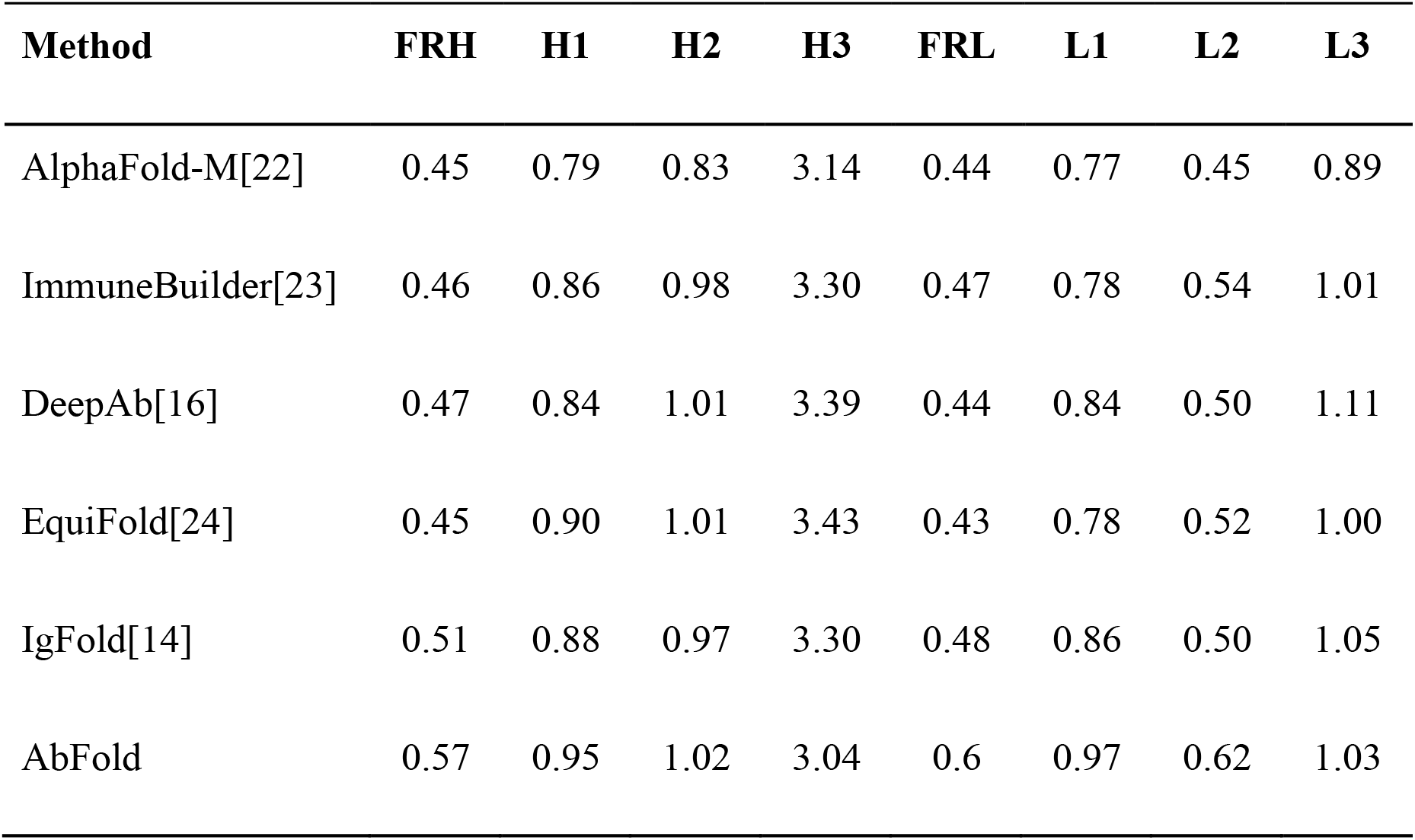
Accuracy comparison of predicted antibody Fv structures

As shown in Table 2, AbFold outperforms other methods in predicting the CDR H3 loop while AlphaFold-Multimer remains state-of-the-art in other regions. A more comprehensive comparison of the five models is presented in Figure 2A. The improvement in the prediction of the CDR H3 loop can be attributed to the modification of low-confidence regions in pre-processed structures.

**Figure 2.**
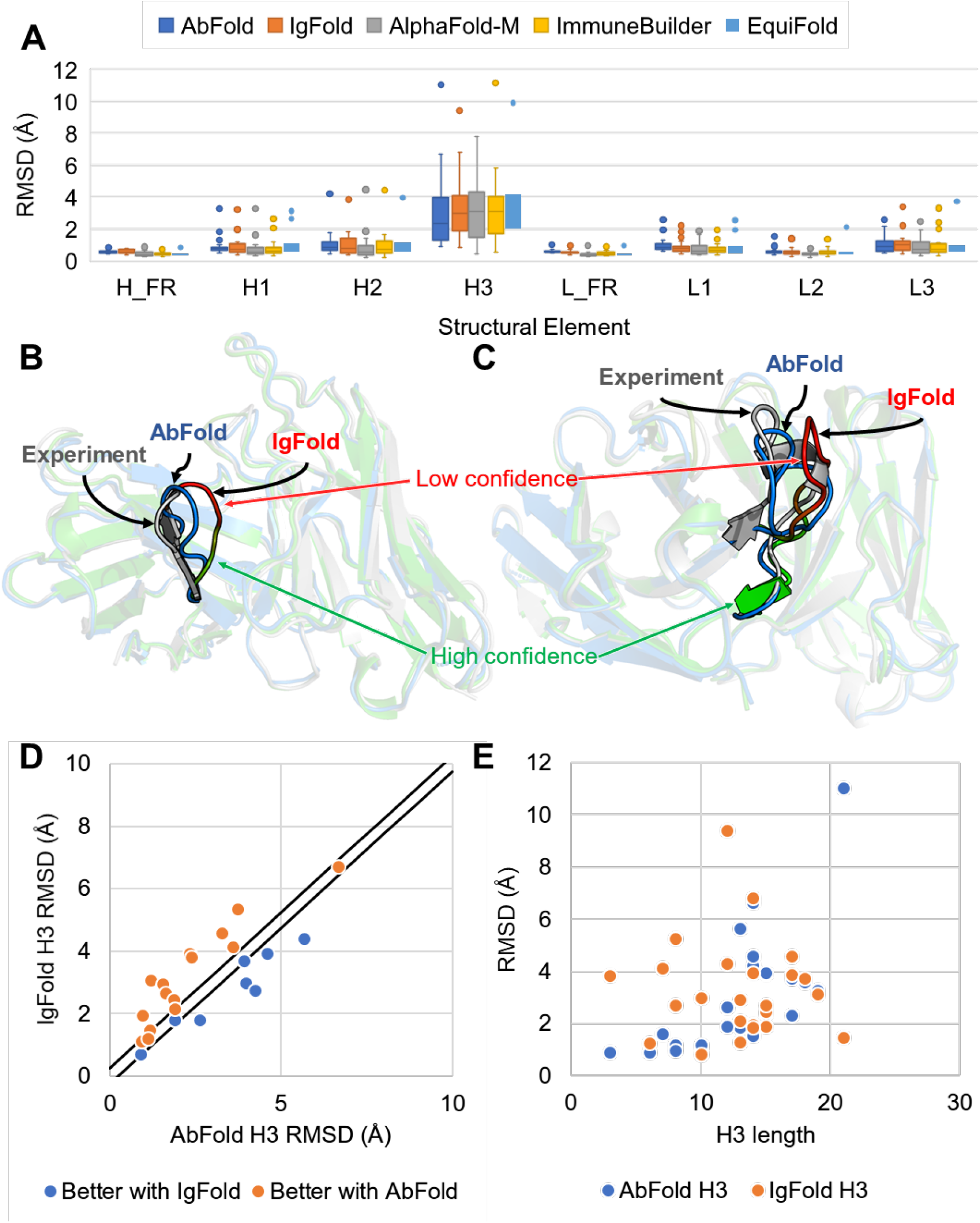
Comparison of structures predicted by different models. (A) Benchmark performance of AbFold, IgFold, AlphaFold-Multimer, ImmuneBuilder, and EquiFold. (B) Comparison of AbFold and IgFold in target 7N0A. Note the predicted structure produced by IgFold is color-coded based on its predicted RMSD, with the regions having a higher predicted RMSD appearing with a richer red coloration. (C) Comparison of AbFold and IgFold in target 7REW. (D) Comparison of CDR H3 RMSD between AbFold and IgFold. (E) Comparison of CDR H3 loop RMSD to the length of CDR H3 loop.

To provide a detailed view of CDR H3 RMSD in each antibody, we compared the results obtained by AbFold and IgFold in a scatter plot (see Figure 2D). The plot indicates that more blue points are concentrated in the top-left region, suggesting that our model exhibits better performance in the majority of antibodies within the benchmark.

Studies have shown that the difficulty of predicting the CDR H3 region is partial driven by the H3 length[16]. To gain a deeper understanding of this phenomenon, we examined the relationship between CDR H3 RMSD and H3 length, as illustrated in Figure 2E. Remarkably, AbFold outperformed IgFold in every H3 length except for one. Additionally, we observed that our model consistently achieved a lower RMSD in shorter H3 regions, which can be attributed to the preference of the point cloud module to extract more information from closer distances.

### Example analysis for CDR H3 Prediction

To gain a deeper understanding of how AbFold optimizes the predicted structure and achieves better accuracy in CDR H3, we analyze two examples from the test sets with PDB ID 7N0A and 7REW. We compare the structures predicted by IgFold and AbFold with the experimental ground truth structure. For 7N0A (as shown in Figure 2B), the CDR H3 loop of this antibody consists of seven residues. IgFold fails to accurately predict the CDR H3 loop conformation (RMSD = 3.07 Å) and exhibits low confidence in this region, whereas AbFold predicts a relatively more correct CDR H3 conformation (RMSD = 1.19 Å). AbFold optimized the orientation of the CDR H3 structure closer to the experimental structure, resulting in a significantly lower RMSD. For 7REW (as shown in Figure 2C), the length of the CDR H3 loop is 14, twice the length of the previous example 7N0A. The conformation predicted by AbFold fits the experimental structure better than IgFold, indicating that AbFold can optimize not only short CDR H3 loops but also longer structures. Notably, in both examples, AbFold’s predictions share a high degree of structural similarity with IgFold’s predictions, except for the CDR H3 loop. These findings support the notion that modifying only the low-confidence regions can lead to improved performance.

Figure 3 compares AlphaFold, IgFold, AbFold, and experimental structures for three additional examples with distinct CDR H3 conformations. AbFold demonstrates superior performance in these cases as well.

**Figure 3.**
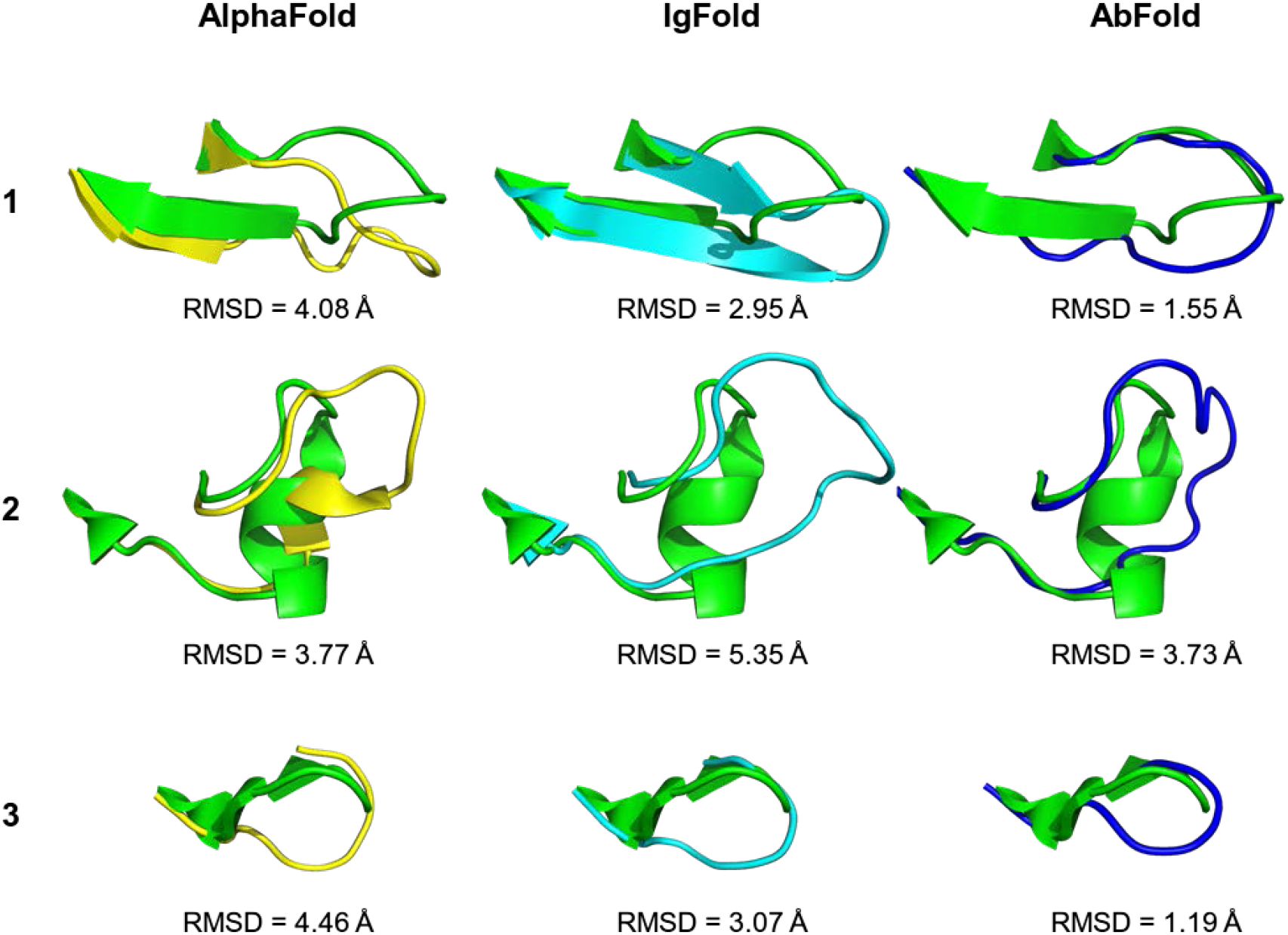
Comparison of CDR H3 loop of predictions from AlphaFold (left, yellow), IgFold (middle, cyan), and AbFold (right, blue) to experimental structure (green). Each row represents the CDR H3 region from one antibody (7mu4, 7zff, 7n0a).

**Figure 4.**
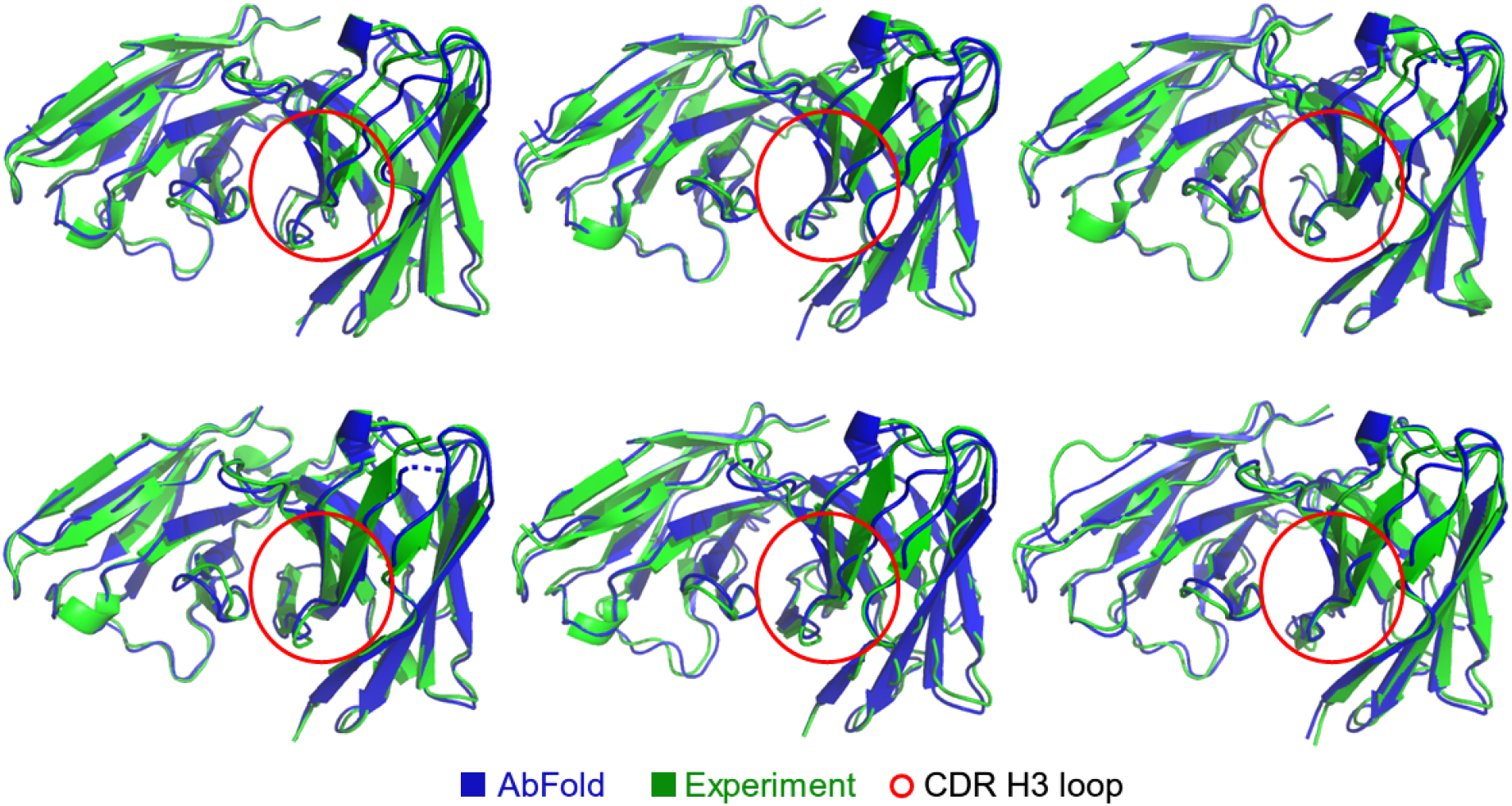
Prediction structure of 6 targets in the benchmark. Blue CDR H3 loops are marked with red circles.

## Discussion

We present AbFold, a new deep-learning structure prediction model for antibodies, that incorporates new techniques such as transfer learning, student-teacher self-supervised learning, point cloud network, and variational autoencoder. We demonstrate that AbFold has a significantly higher accuracy in the CDR H3 region and achieves a comparable result in other regions.

AlphaFold2 demonstrates great capability in predicting general protein 3D structure, we use the Evoformer block in AlphaFold to extract and transfer pre-trained knowledge.

We also adopt a teacher-student model to make use of a large amount of unlabeled antibody data.

Previous methods usually focus on sequences data, using either MSA or language model to capture patterns inside the sequence. Considering the CDR H3 region in antibody structures are highly flexible, AbFold incorporates a Point-MAE module and a VAE module. These two modules focus on 3D atom interactions and are proven to improve accuracy in CDR H3 regions.

We have found certain limitations while designing and testing out AbFold: both training and test datasets are relatively limited in size, which may make the model comparison not as robust; the dynamism of CDR loops may be better modeled with specific antigen pair, instead of on a stand-alone basis.

## Acknowledgment

This work was partly supported by National Natural Science Foundation of China (No.62106051) and Shanghai Pujiang Program (No.21PJ1400600).

