## Supplementary information for "AbFold -- an AlphaFold Based Transfer Learning Model for Accurate Antibody Structure Prediction"

### *Ablation Study*

In order to identify the key design elements of our proposed AbFold model, we carried out experiments on ablated versions that lacked point cloud, variational autoencoder, or semi-supervised training components. Moreover, we investigated the effect of different loss functions.

**Model structures.** Table S1 reports the outcomes of our experiments in which we removed different parts of the model structures. We found that the absence of any component led to varying degrees of performance degradation on both the framework regions and CDR H3 loop, indicating that all three components are crucial for accurately predicting antibody structures. Moreover, we observed that the model lacking variational autoencoder performed poorly in predicting the H chain structure, as depicted in Table S1. In terms of the CDR H3 loop, Figure S1 displays the RMSD distribution of the structures predicted by each model, revealing a significant difference between the model without variational encoder and the one without semi-supervised training. This suggests that the variational encoder can effectively model highly variable antibody regions. Furthermore, our experiments demonstrated that the model's overall performance was inferior without the semi-supervised training than with it, highlighting the pivotal role of this training in our approach.

| **Model** | **Fr H** | **Fr L** | **H3** |
|---|---|---|---|
| Without point cloud | 0.94 | 0.83 | 3.50 |
| Without variational autoencoder | 2.62 | 0.74 | 5.65 |
| Without semi-supervised training | 0.91 | 0.70 | 3.73 |
| AbFold | 0.72 | 0.71 | 3.07 |

Table S1: Ablation study on model structures. We compare the RMSD calculated over backbone heavy atoms of each method with unit Å.

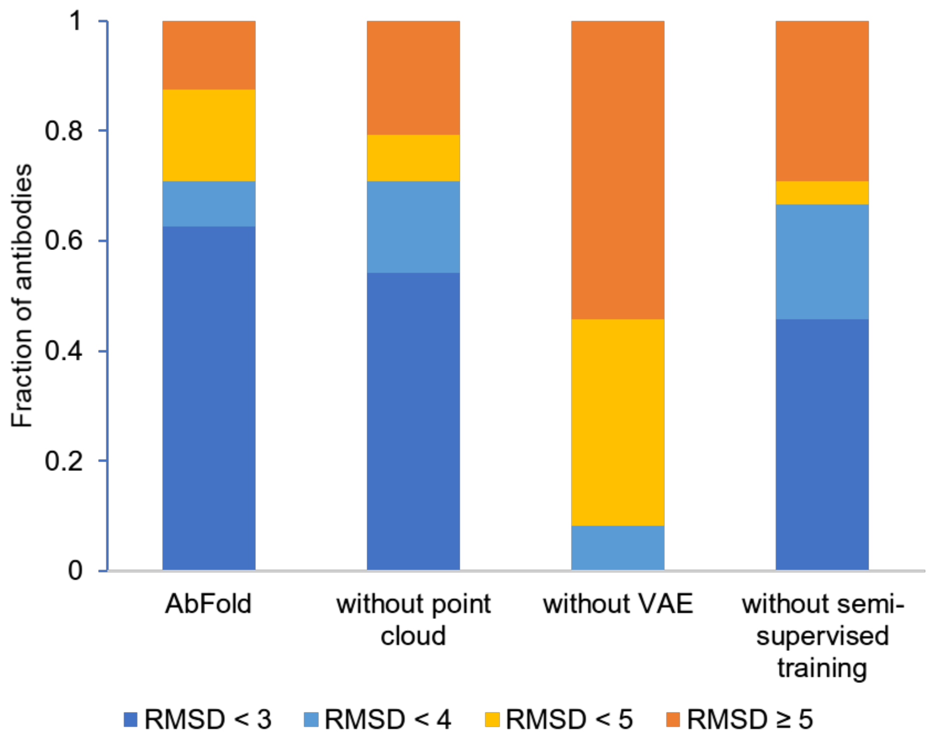


Figure S1. RMSD distribution of CDR H3 loops predicted by each model. The model without variational autoencoder displays much worse RMSD distribution compared to the other models.

**Loss functions.** To explore the potential effects of different loss functions, we integrated several functions from AlphaFold2 and IgFold into our network. Initially, we used FAPE and Structure Violation as our targets for supervision, but we observed that some bonds between residues were longer than expected, resulting in suboptimal results. To address this issue, we added bond length loss to our loss functions in subsequent experiments, which successfully resolved the problem of unreasonable bond length. Additionally, we attempted to incorporate supervised chi loss to enhance side chain structure optimization, but it had an adverse effect on backbone structure and was therefore excluded from our training process. To promote training stability, we implemented a two-stage training approach, each utilizing different loss functions. The first stage exclusively employed FAPE loss, while the second stage incorporated both supervised chi loss and bond length loss as targets. This strategy effectively led to convergent results while avoiding unreasonable structures.
